# Comparison of Vitek^®^ 2, three different gradient strip tests and broth microdilution for detecting *vanB*-positive *Enterococcus faecium* isolates with low vancomycin MICs

**DOI:** 10.1101/634535

**Authors:** Ingo Klare, Jennifer K. Bender, Carola Fleige, Nancy Kriebel, Axel Hamprecht, Sören Gatermann, Guido Werner

**Affiliations:** National Reference Centre for Staphylococci and Enterococci (NRC), Division Nosocomial Pathogens and Antibiotic Resistances, Department of Infectious Diseases, Robert Koch Institute, Wernigerode Branch, Wernigerode, Germany; Institute for Medical Microbiology, Immunology and Hygiene, University of Cologne, Cologne, Germany and DZIF (German Centre for Infection Research), partner site Bonn-Cologne, Germany; Department of Medical Microbiology, Institute for Medical Microbiology and Hygiene, Ruhr University Bochum, Bochum, Germany

**Keywords:** EUCAST, VRE diagnostic, vancomycin resistance, *vanB*

## Abstract

**Objectives:** In 2018, EUCAST issued a warning regarding unreliable results of gradient strip tests for confirming vancomycin resistance in enterococci. We compared the performance of various diagnostic standard and confirmatory tests to identify and determine *vanB*-type vancomycin resistance.

**Methods:** We analysed a collection of *vanB*-positive *E. faecium* isolates (n = 68) with low level vancomycin minimal inhibitory concentrations (MICs) and compared the performance of Vitek^®^ 2 (bioMérieux), of broth microdilution and of three different gradient strip test providers (Oxoid, Liofilchem, bioMérieux). For the latter we compared the standard procedure vs. a protocol with increased inoculum, a rich agar medium and a longer incubation time (“macromethod”).

**Results:** The sensitivity of Vitek^®^ 2 was 81% of *vanB*-isolates compared to 72% for broth microdilution and 61 – 63% for the three gradient strip tests using standard conditions. The “macromethod” substantially improved performance of all strip tests resulting in a sensitivity of 89 – 96% at 48h readout.

**Conclusions:** The “macromethod” provided the overall best performance for recognition of *vanB E. faecium*. We therefore propose to adapt the EUCAST warning and to recommend the “macromethod” with an additional 48h readout or a confirmation by a *vanB*-specific PCR from culture.

## Introduction

Many hospitals in Germany and other European countries are challenged by an increasing number of vancomycin-resistant enterococci (VRE) associated with colonizations and infections in hospitalized patients. The rising numbers of VRE are mainly driven by an increase in *vanB*-positive VRE locally^1^ or on a country-wide scale.^2^ Furthermore, there is a growing recognition of *vanB*-VRE with low level vancomycin MICs just below the breakpoint of 4 mg/L.^3,4^ Uncertain diagnostic results may rely on confirmation by alternative tests such as MIC gradient strip assays (e.g., Etest). The EUCAST issued a warning in July 2018 regarding less reliable strip assay results for determining and confirming vancomycin resistance in enterococci (http://www.eucast.org/ast_of_bacteria/warnings/) leaving the diagnostic community with an uncertainty how to confirm *vanB-*positive VRE with low level vancomycin resistance. For the present study, we established a strain collection of pre-characterized and heterogeneous *vanB*-positive *E. faecium* isolates from all over Germany (n = 80) and from recent years which all had low level vancomycin MICs in previous standard diagnostic assays and/or ambiguous resistance phenotypes. We aimed at comparing the performance of five diagnostic standard and confirmatory tests to identify and determine *vanB*-type vancomycin resistance in the most reliable manner.

## Materials and Methods

The 68 *vanB*-positive *E. faecium* isolates originated from all over Germany and displayed low level vancomycin MICs in previous diagnostic assays (see **Supplementary Table S1**). The collection was especially enriched with isolates demonstrating vancomycin MICs of 2 to 4 mg/L (S) and 8 mg/L (R) in broth microdilution. We did not include *vanB* strains with vancomycin MICs of ≤1 mg/L as these strains may possess defects in *vanB* regulation.^5^ To exclude any bias in the strain collection, we included isolates from 42 diagnostic laboratories submitted between 2015 and 2018 to the National Reference Centre. We determined vancomycin MICs using Vitek^®^ 2 card AST P611 (bioMérieux, Nurtingen, Germany), broth microdilution (according to EUCAST standards) and gradient strip assays from three providers (M.I.C. Evaluator™, Oxoid/Thermo Fisher Scientific, Wesel, Germany; MIC test strip^®^, Liofilchem, Roseto degli Abruzzi, Italy; Etest^®^, bioMérieux). For gradient strip assays, we compared the standard procedure vs. the “macromethod”, which includes a rich agar medium (Brain Heart Infusion instead of Mueller-Hinton), a higher inoculum (McFarland 2 instead of 0.5) and a longer incubation time of up to 48h. *E. faecalis* ATCC20912, *E. faecium* ATCC19434, *E. gallinarum* BM4174 (*vanC1*) and *E. casseliflavus* ATCC25788 (*vanC2)* were used as reference isolates. Ten vancomycin-susceptible *E. faecium* (negative for *vanA* and *vanB*) served as a control group. Presence of *vanB* in the study group isolates and absence in control and reference isolates was confirmed by PCR.^4^ Statistical calculations for sensitivity and specificity were carried out according to: https://www.medcalc.org/calc/diagnostic_test.php.

## Results

The ten vancomycin-susceptible *E. faecium* of the control group and the susceptible reference isolates all revealed MICs below 4 mg/L in all assays and under all conditions demonstrating a specificity of 100% (95% CI: 69-100).

Vitek 2 identified 52 of the 68 *vanB*-VRE (sensitivity 81% [95% CI: 71 – 89]; **Table**). Using broth microdilution, only 41 *vanB*-VRE were identified after 24h demonstrating a sensitivity of 72% (95% CI: 61 – 80) for the “Gold standard”. Incubation of the plates for another 24h allowed the identification of all 68 *vanB*-VRE (MICs of >4 mg/L).

The comparison of three gradient strip assays revealed similar results for the vancomycin Etest, vancomycin MIC test strip and vancomycin M.I.C. Evaluator: altogether 26, 24 and 28 *vanB*-VRE were correctly identified leading to sensitivities of 61 – 63% (**Table, Figure**; see also **Supplementary Figures S1** and **S2**). Sensitivities were slightly higher after 48h readout (65 – 69 %; **Table**).

Using the “macromethod” (see Methods), as explicitly suggested by bioMérieux for their Etest, substantially improved the sensitivity of all gradient strip assays. After overnight incubation 57 *vanB*-VRE were identified by Etest (sensitivity 86% [CI: 76 – 93]), 45 by MIC test strip (sensitivity 75% [CI: 65 – 83]) and 58 by M.I.C. Evaluator (sensitivity 87% [CI: 77 – 94]). Sensitivity further improved after 48h readout with 63 *vanB*-VRE (93% [CI 85 – 98]) for Etest, 60 (89% [CI: 80 – 95]) for MIC test strip and 65 (96% [CI: 88 – 99]) for M.I.C. Evaluator.

## Discussion

Since their first description in the early 1990s, low level vancomycin MICs were predominantely reported for *vanB*-type enterococci compared to *vanA*-type VRE. The phenomenon is most probably caused by different two-component circuits mediated via VanR/VanS and VanR_B_/VanS_B_ regulating induction of vancomycin resistance in *vanA*-type and *vanB*-type VRE, respectively. Susceptibility test norm and standard providers like CLSI and EUCAST recognized this by broadening the intermediate category for vancomycin (CLSI) or lowering the vancomycin clinical breakpoint (EUCAST). However, correct identification especially of *vanB*-type VRE remains challenging.^4–6^ The sensitivity for detection of *vanB*-type VRE was 81% for Vitek 2 compared to 72% for broth microdilution and 61-63% for the three gradient strip tests. The “macromethod” substantially improved sensitivities of all strip tests to 89–96% after 48h readout. Specificity remained excellent when using the “macromethod” (100% [95% CI: 69 – 100]).

Previous attempts to improve detection of *vanB*-type VRE included a supplementation of 10 g/L oxgall to agar media (Mueller-Hinton, Brain Heart Infusion). This increases *vabB* cluster gene expression and thereby allowed improved detection of *vanB*-VRE with low vancomycin MICs.^3^ Unfortunately, media with this supplement are not commercially available. Applying the “macromethod” with commercial kits and media might be implemented much easier and would simply require an extended incubation of up to 48h. We are well aware that an additional working day is in conflict with demands for quick and reliable diagnostics and for decisions for infection control, for instance, in cases of admission screenings. Nevertheless, using the “macromethod” with 24h readout was already superior to the “Gold standard” broth microdilution. PCR-based screening directly from clinical samples may have certain advantages in identifying *vanB*-type resistance, but the reservoir and frequent occurrence of *vanB* in human intestinal commensal bacteria conflicts with a reliable test result and inevitably demands a confirmation by culture.^7,8^

In EUCAST expert rules v3.2 which are currently (04/2019) in a wide consultation process, it is recommended that enterococci with a positive *vanB* result which appear vancomycin-susceptible should be reported resistant to vancomycin (http://www.eucast.org/documents/consultations/). We fully support this deduction which is in line with our study results demonstrating that sensitivity of currently available methods needs improvement and minor methodological changes may influence the categorization “susceptible or resistant to vancomycin.

In conclusion, we recommend to change the present EUCAST warning against the general use of MIC strips and to specify that the “macromethod” should be employed for confirming vancomycin resistance in *Enterococcus* spp. Strip providers not recommending this method should adapt their application guidelines. For clinically relevant enterococci where a rapid therapeutic decision is needed, e.g. for isolates from blood stream infections, a molecular test (e.g., PCR) can be recommended in order to save time and to further increase sensitivity.

## Transparency Declaration and Funding

This work was supported by a Grant of the Federal Ministry of Health, Germany, to the work of the National Reference Centre for Staphylococci and Enterococci. The study was performed under the auspices of the Section “Basics” of the Paul Ehrlich Society for Chemotherapy and the German National Antibiotic Susceptibility Test Committee NAC (www.nak-deutschland.org).

**Table.** Performance of different primary and confirmatory diagnostic assays in detecting *vanB*-mediated vancomycin resistance in *E. faecium* isolates (n = 68). Ten vancomycin-susceptible and *vanB*-negative *E. faecium* isolates served as a control group.

| Susceptibility test method | Test version | VRE detected N = | Sensitivity in % | Sensitivity (95% CI) |
| --- | --- | --- | --- | --- |
| n.a. | ***vanB*-PCR (reference)** | **68** | **100** | **(95 – 100)** |
| **Broth microdilution** | 24h | **41** | **72** | **(61 – 80)** |
|  | 48h | **68** | **100** | **(95 – 100)** |
| **bioMérieux VITEK® 2** | 20h | **52** | **81** | **(71 – 89)** |
| **bioMérieux Etest®** | Standard (24h) | **26** | **62** | **(52 – 71)** |
|  | Standard (48h) | **38** | **69** | **(59 – 78)** |
|  | Macromethod (24h) | **57** | **86** | **(76 – 93)** |
|  | Macromethod (48h) | **63** | **93** | **(85 – 98)** |
| **Liofilchem MIC test strip®** | Standard (24h) | **24** | **61** | **(51 – 70)** |
|  | Standard (48h) | **32** | **65** | **(55 – 74)** |
|  | Macromethod (24h) | **45** | **75** | **(65 – 83)** |
|  | Macromethod (48h) | **60** | **89** | **(80 – 95)** |
| **OXOID M.I.C. Evaluator®** | Standard (24h) | **28** | **63** | **(53 – 72)** |
|  | Standard (48h) | **36** | **68** | **(58 – 77)** |
|  | Macromethod (24h) | **58** | **87** | **(77 – 94)** |
|  | Macromethod (48h) | **65** | **96** | **(88 – 99)** |

**Table.** Performance of different primary and confirmatory diagnostic assays in detecting vαnS-mediated vancomycin resistance in *E. faecium* isolates (n = 68). Ten vancomycin-susceptible and *vonB*-negative *E. faecium* isolates served as a control group.

| Susceptibility test method | Test version | VRE detected<br>N = | Sensitivity<br>in % | Sensitivity<br>(95% CI) |
| --- | --- | --- | --- | --- |
| n.a. | ***vanB*-PCR (reference)** | **68** | **100** | **(95 – 100)** |
| **Broth microdilution** | 24h | **41** | **72** | **(61 – 80)** |
|  | 48h | **68** | **100** | **(95 – 100)** |
| **bioMérieux VITEK® 2** | 20h | **52** | **81** | **(71 – 89)** |
| **bioMérieux Etest®** | Standard (24h) | **26** | **62** | **(52 – 71)** |
|  | Standard (48h) | **38** | **69** | **(59 – 78)** |
|  | Macromethod (24h) | **57** | **86** | **(76 – 93)** |
|  | Macromethod (48h) | **63** | **93** | **(85 – 98)** |
| **Liofilchem MIC test strip®** | Standard (24h) | **24** | **61** | **(51 – 70)** |
|  | Standard (48h) | **32** | **65** | **(55 – 74)** |
|  | Macromethod (24h) | **45** | **75** | **(65 – 83)** |
|  | Macromethod (48h) | **60** | **89** | **(80 – 95)** |
| **OXOID M.I.C. Evaluator®** | Standard (24h) | **28** | **63** | **(53 – 72)** |
|  | Standard (48h) | **36** | **68** | **(58 – 77)** |
|  | Macromethod (24h) | **58** | **87** | **(77 – 94)** |
|  | Macromethod (48h) | **65** | **96** | **(88 – 99)** |

**Figure.**
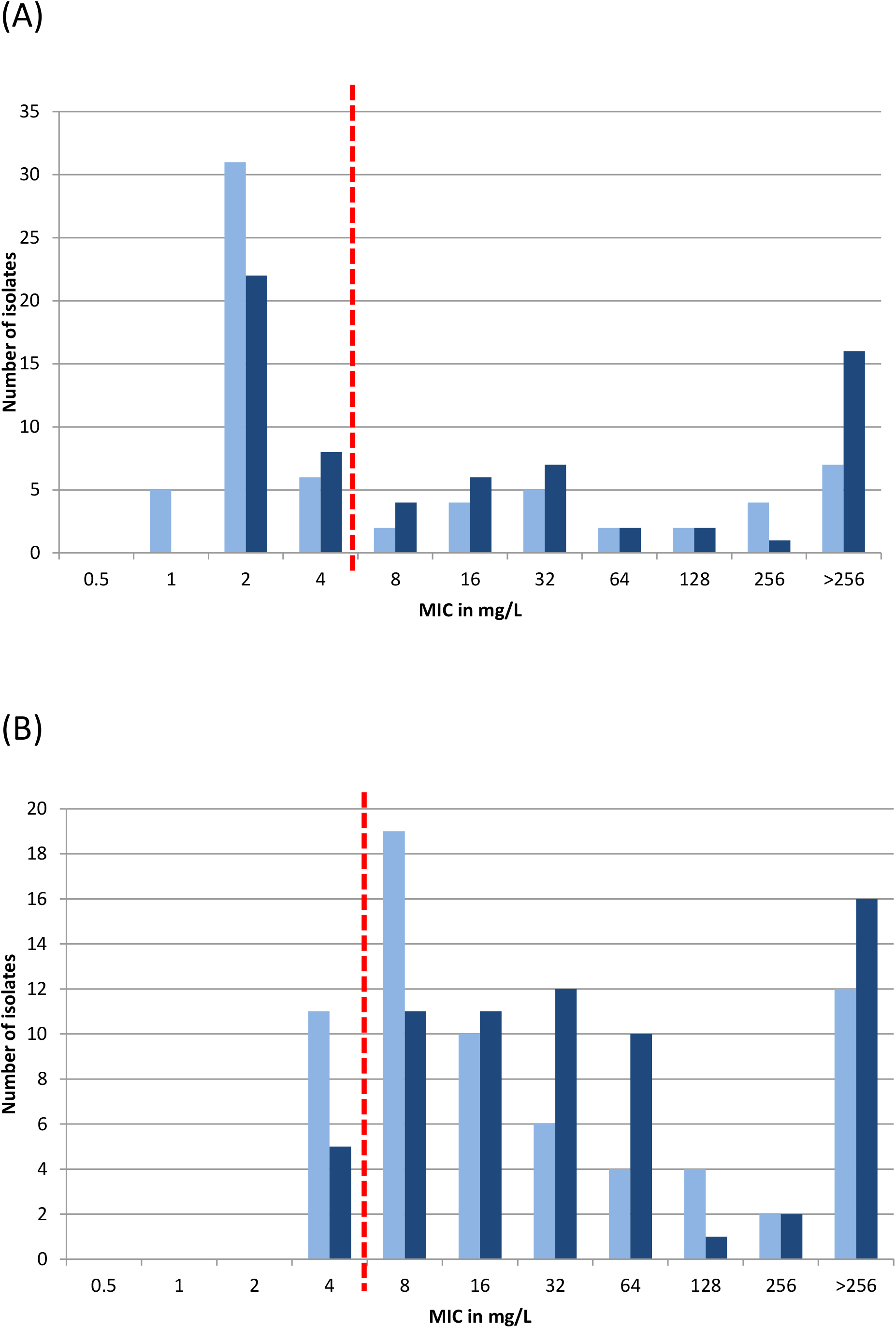
Performance of the Etest^®^ (Biomerieux) by the standard method (A) vs. the “macromethod” (B) for detection of *vonB*-positive *E. faecium* (n = 68). The red dotted line corresponds to the EUCAST clinical breakpoint for vancomycin (R >4 mg/L). Light blue columns represent MIC results after 24h, dark blue columns after 48h readout.

## Acknowledgements

We thank Christine Günther for excellent technical assistance. We are grateful to all clinical diagnostic laboratories in Germany sending strains for further analysis and typing to the National Reference Centre for Staphylococci and Enterococci.

## Supplementary Figures

**Figure S1. Performance of the Liofilchem MIC test strip^®^ by the standard method (A) vs. the “macromethod” (B) for detection of *vanB*-positive *E. faecium* (n = 68).** The red dotted line corresponds to the EUCAST clinical breakpoint for vancomycin (R >4 mg/L). Light blue columns represent MIC results after 24h, dark blue columns after 48h readout.

**Figure S2. Performance of the OXOID M.I.C. Evaluator™ gradient strip by the standard method (A) vs. the “macromethod” (B) for detection of *vanB*-positive *E. faecium* (n = 68).** The red dotted line corresponds to the EUCAST clinical breakpoint for vancomycin (R >4 mg/L). Light blue columns represent MIC results after 24h, dark blue columns after 48h readout.

**Table S1. Enterococcal isolates included in this study**. Data display the origin of isolates, the results of vancomycin broth microdilution, vancomycin Vitek 2 analyses and the three vancomycin gradient strip tests; the latter results are further specified depending on the method used, standard method vs. the “macromethod”, and subsequent 24h/48h incubation. MIC results of strips providing odd concentrations which do not represent doubling dilutions such as 0.75 or 1.5 mg/L (Etest, MIC test strip) were extrapolated to even concentrations (e.g., 0.75 to 1 mg/L or 1.5 to 2 mg/L).

