## Supplementary Figure S2 for "Comparison of Vitek^®^ 2, three different gradient strip tests and broth microdilution for detecting *vanB*-positive *Enterococcus faecium* isolates with low vancomycin MICs"

(A)

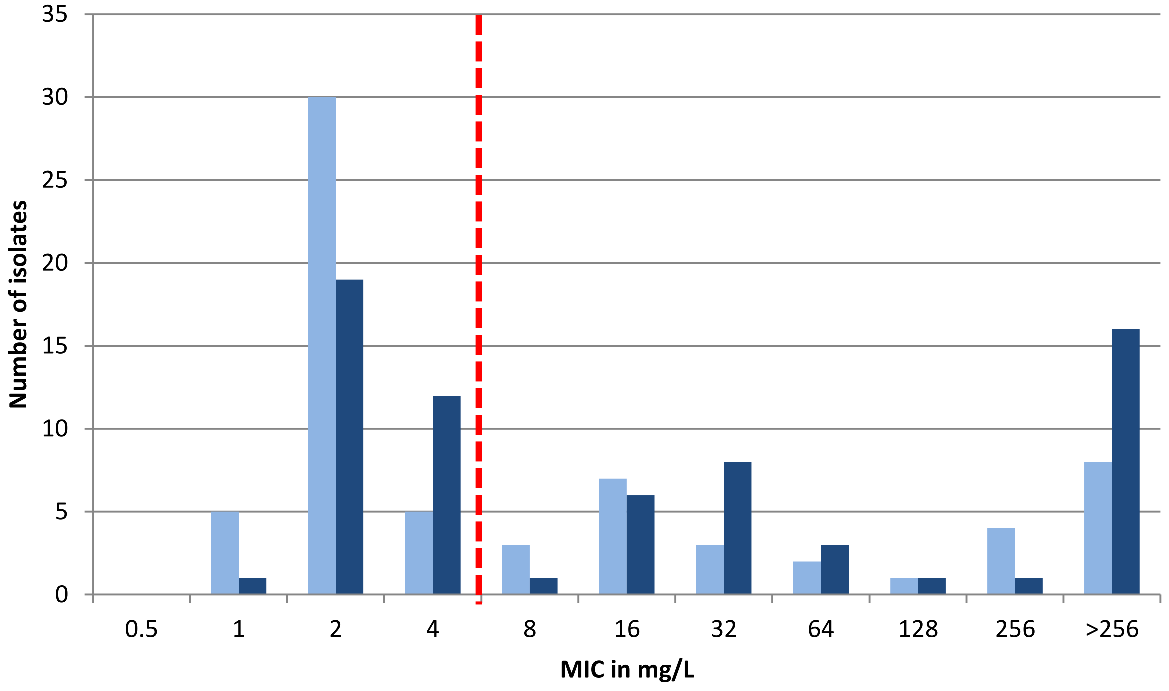


(B)

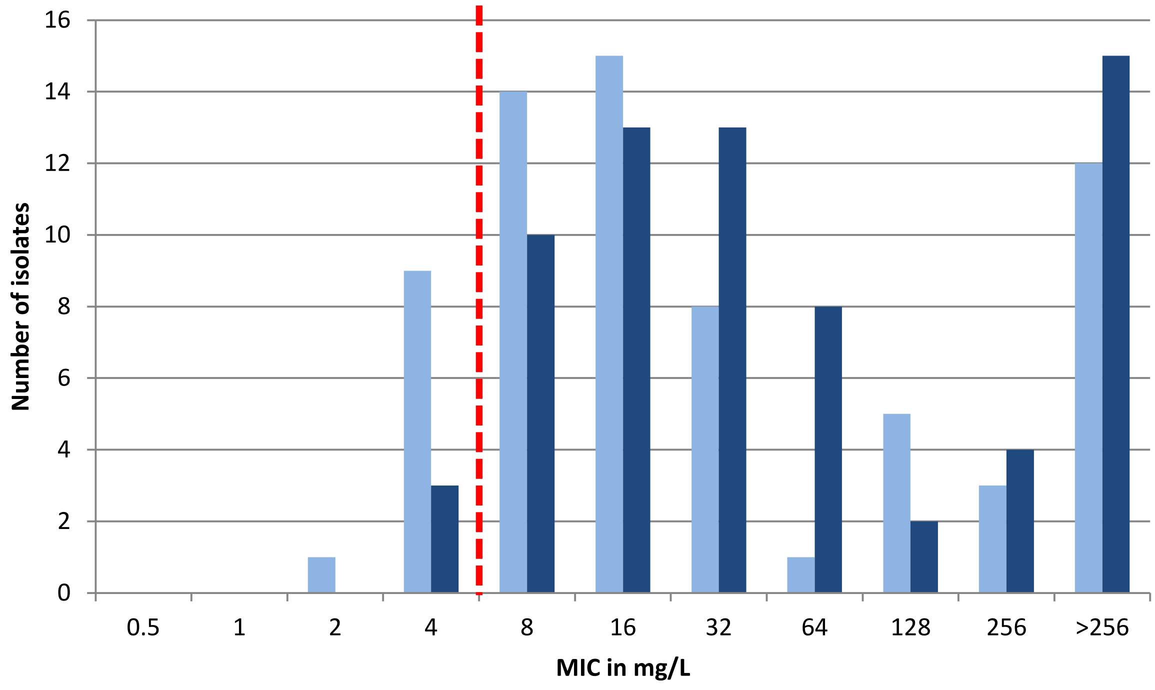


**Supplementary Figure S2. Performance of the OXOID M.I.C. Evaluator™ gradient strip by the standard method (A) vs. the "macromethod" (B) for detection of *vanB*-positive *E. faecium* (n = 68).** The red dotted line corresponds to the EUCAST clinical breakpoint for vancomycin (R >4 mg/L). Light blue columns represent MIC results after 24h, dark blue columns after 48h readout. (see main manuscript for further details).
